# Oncometabolite fumarate impairs ATR-CHK1 signaling by succinating RPA1 in Fumarate Hydratase-deficient renal cell carcinoma cells

**DOI:** 10.64898/2025.12.08.692638

**Authors:** Nian Liu, Changxian Shen, Tingting Zhou, Yu Zhou, Hang Yuan, Hongjie Bi, Yingying Wang, Xinyu Pei, Rui Su, Li Zheng, Yihui Shi, Binghui Shen

**Affiliations:** Department of Cancer Genetics and Epigenetics, Beckman Research Institute, City of Hope, 1500 East Duarte Road, Duarte, California, CA91010, USA; Division of Surgical Oncology, Department of Surgery, Ohio State University Comprehensive Cancer Center and Ohio State University Wexner Medical Center, Columbus, OH43210, USA; Department of Systems Biology, Beckman Research Institute of City of Hope, Monrovia, CA91016, USA; California Pacific Medical Center Research Institute, Sutter Bay Hospitals, San Francisco, CA94107, USA

**Keywords:** Fumarate hydratase-deficient renal cell carcinoma (FHdRCC), Replication Protein A1 (RPA1), Succination, ATR-CHK1 pathway, genome instability

## Abstract

Abnormal accumulation of oncometabolite fumarate drives susceptibility in fumarate hydratase-deficient renal cell carcinoma (FH-dRCC), but the precise mechanisms remain not fully understood. In this study, we demonstrate that high fumarate levels impair activation of ATR-CHK1 signaling in response to replication stress and DNA damage. Mechanistically, fumarate modifies RPA1, an essential factor for ATR-CHK1 activation through succination, a post-translational modification. Succination of RPA1 occurs mainly at cysteine residues 481 and 486, which reduces its binding affinity for single-stranded DNA (ssDNA). RPA1 succination leads to deficient recruitment of TOPBP1 to ssDNA, resulting in attenuated CHK1 activation and defective cell cycle arrest in response to DNA damage. Succinated RPA1 compromises homologous recombination-mediated DNA repair. Our findings establish that fumarate-induced succination of RPA1 impairs DNA repair and cell cycle control, promoting genomic instability in FH-dRCC. This work reveals a novel mechanism by which oncometabolites contribute to genomic instability.

## 1. Introduction

Fumarate hydratase (FH) is a crucial enzyme in the tricarboxylic acid cycle (TCA), and its loss is implicated in various cancers, notably in renal cell carcinoma (RCC) [1, 2]. FH-deficient RCC (FH-dRCC) is caused by germline mutations of FH and characterized by abnormal accumulation of fumarate, leading to metabolic reprogramming, epigenetic changes, and altered cellular signaling pathways [3–6]. Fumarate, succinate and 2-HG are well-established oncometabolites, which promote tumorigenesis via multiple mechanisms, especially epigenetic regulation of gene expression and chromatin remodeling; distinct from succinate and 2-HG, fumarate can covalently react with the thiol group of cysteines, giving rise to fumarate–cysteine adducts in a chemical reaction referred to as succination [7]. Succination affects multiple proteins in FH-deficient cells and their microenvironment [8]. For example, KEAP1 succination leads to the stabilization and nuclear translocation of NRF2, which in turn induces the expression of genes involved in antioxidant response elements (AREs) and activates a strong antioxidant response [9]. In the nucleus, succination of SMARCC1 causes depletion of the tumor-suppressor gene SNF5, resulting in alteration of chromatin remodeling [10]. In the FH^-/-^ tumor microenvironment, the accumulated fumarate can induce ZAP70 succination which decreases T cell activation [11]. Fumarate can also regulate B-cell activation and function by directly succinating and inactivating LYN [12]. Fumarate-mediated PTEN succination leads to PTEN inactivation and consequently activation of the PI3K-AKT axis to promote FH-deficient tumor cell proliferation [13]. Thus, mounting or increasing evidence has highlighted the pivotal roles of succination in the various processes of cell physiology.

Genomic instability is a hallmark of cancer cells [14]. In hereditary cancers, genomic instability results from mutations in DNA repair genes and drives cancer development [15]. The ATR-CHK1 pathway plays the principal role in maintaining genome stability, especially in response to DNA replication stress and damage [16]. ATR-CHK1 signaling maintains genome stability via many mechanisms, including arresting cell cycle at S and G2/M phases, promoting DNA repair, stabilizing stalled replication forks, increasing dNTPs for DNA repair and replication, and promoting DNA replication restart [17]. ATR-CHK1 signaling can be activated by both endogenous and exogenous genotoxins, such as stalled replication forks, DNA repair intermediates, R-loop, and aberrant telomeres as well as chemotherapy and radiotherapy agents-induced single-strand DNA (ssDNA) and DNA double strand breaks (DSBs) [18]. Molecularly, ATR-CHK1 is activated by replication protein A (RPA)-coated ssDNA. RPA1 is the largest subunit of the RPA complex, which is involved in ssDNA and DSBs repair pathways. RPA1 is essential for cell survival and maintenance of chromosomal stability [19]. RPA-coated ssDNA is the signal and platform for initiating ATR-CHK1 checkpoint pathway activation by recruiting numerous proteins including ATR, ATRIP, and TOPBP1 [20, 21]. The functions of RPA1 protein can be regulated by multiple post-translational modifications in response to genotoxic stress to dynamically regulate DNA repair processes, including acetylation, ubiquitination, SUMOylation [22–25].

Metabolic reprogramming is another hallmark of cancer cells [26]. However, the functional linkage and mechanisms by which metabolic reprogramming impacts on cancer genome instability remain enigmatic. In this study, we demonstrate that succination of RPA1 significantly reduces its ability to bind ssDNA, which impairs the subsequent recruitment of the ATR-ATRIP complex and the essential co-activator TOPBP1. This disruption leads to impaired activation of the ATR-CHK1 checkpoint pathway, compromising the cell’s ability to respond effectively to replication stress and DNA damage. Our findings reveal a direct link between metabolic dysregulation and genome instability through succination of the key DNA repair proteins.

## 2. Materials and Methods

### 2.1. Antibodies

Anti-pCHK1-S345 (2348S; 1:1000), anti-CHK1 (2360T;1:1000), anti-pCHK2-T68 (2661;1:1000), anti-CHK2 (6334T,1:1000), anti-DYKDDDDK (8146;1:1000), anti-ATRIP (2737;1:1000), anti-ATR (2790;1:1000), anti-TOPBP1 (14342;1:1000), anti-RPA70 (2267; western-blot dilution:1:1000; immunostaining dilution 1:500); anti-Tubulin (2144;1:1000), anti-Histone H3 (4499;1:1000), anti-GAPDH (2118;1:1000) were purchase from cell signaling. Anti-FH (SC-393992; 1:500) was purchased from Santa Cruz Biotechnology and anti-2SC (CRB2005017_3; 1:500) was purchased from Biosynth. Anti-rabbit IgG (7074; 1:2000) and anti-mouse IgG (7076; 1:2000) HRP-linked antibodies were purchased from ThermoFisher Scientific.

### 2.2. Chemicals

Hydroxyurea (HY-B0313), fumarate hydratase inhibitor FH-IN-1(HY-100004) were purchased from MedChemExpress. Dimethyl fumarate (DMF) was purchased from Sigma-Aldrich. Sodium fumarate (215530050) was purchased from Thermo Scientific.

### 2.3. Cell lines

HK-2 cells and HEK293T cells were obtained from ATCC and cultured in DMEM medium with 10% fetal bovine serum (FBS) and 1% penicillin-streptomycin antibiotics (Thermo Fisher Scientific).786-0 cells were obtained from ATCC and cultured in RPMI1640 medium with 10% FBS and 1% penicillin-streptomycin antibiotics. UOK262 and UOK268 cells were from W. M. Linehan, NCI Urologic Oncology Branch[27]. UOK262 and UOK268 cells, UOK262 and UOK268 cells stably expressing Flag-tagged FH were cultured in DMEM medium with 10% FBS, 1% NEAA and 1% penicillin-streptomycin antibiotics. Cells were maintained in a humidified incubator at 37°C with 5% CO2.

### 2.4. Plasmid construction, mutagenesis and transfection

For FH overexpression in UOK262 and 268 cells, polymerase chain reaction (PCR)-amplified human wild type FH were cloned to pSin4-Flag lentivirus vector. Lentiviral particles were generated by transient transfection of HEK293T cells with pSin4-FH vector, the packaging plasmid psPAX2 and the envelope plasmid pMD2.G using lipofectamine 3000 according to the manufacturer’s instructions. Viral supernatants were collected at 48h and 72h post-transfection. UOK262 and UOK268 cells were plated at 60% confluence and transduced by lentivirus with polybrene (8µg/mL). At 48 h post-infection, cells were selected with puromycin (2µg/mL) for 5 days, and then transduction efficiency was verified by Western blotting.

Human pCMV-RPA1-flag expression plasmid was synthesized by Sino Biological Inc. Point mutations of RPA1-flag expression plasmid were constructed using a site-directed mutagenesis approach. Briefly, complementary primers containing the desired nucleotide substitution were used to amplify the plasmid by PCR. The parental DNA was digested with DpnI, and then PCR product was transformed into DH5-α cells, and colonies were screened by Sanger sequencing to confirm mutation. Cells were transfected with plasmid DNA using lipofectamine 3000 according to the manufacturer’s protocol.

### 2.5. LC-MS/MS

To identify the succinated protein in UOK268 cells, total cell lysates extracted from UOK268 cells were pulled down by 2-SC antibody conjugated to magnetic anti-flag beads (normal IgG as negative control), then eluted with 3×Flag peptide. The samples were processed by LC-MS/MS analysis. For identification of the succination modification sites, purified recombinant RPA1 protein was incubated with 100 mM sodium fumarate in the buffer (50 mM Tris-HCl pH 7.5) at 37 °C for 2 h. Control was performed in the absence of sodium fumarate. The samples were subjected to SDS-PAGE. Proteins were visualized by Coomassie staining. Gel bands corresponding to RPA1 were excised and analyzed by LC–MS/MS.

### 2.6. Immunoprecipitation and immunoblotting

For immunoprecipitation (IP), cells were lysed in a buffer containing 1% Nonidet P-40, 50 mM Tris–HCl, 0.1 mM EDTA, 150 mM NaCl, and proteinase inhibitors. Whole cell lysates were precipitated with Pierce^TM^ Protein A/G Magnetic Beads (ThermoFisher Scientific). The proteins were eluted with 2×SDS loading buffer and analyzed by western blot following the standard protocol. For immunoblotting, the cultured cells were lysed in RIPA lysis buffer supplemented with phosphatase and protease inhibitors cocktails (Sigma). The proteins were separated by SDS-PAGE and transferred to nitrocellulose membrane The membranes were blocked in a 5% non-fat milk for 1h at 37°C and were incubated with primary antibodies against specific proteins of interest at 4°C overnight, followed by incubation with HRP-conjugated secondary antibodies.

### 2.7. Cell cycle analysis

Cells were seeded at density of 2 × 10⁵ cells/well in 6-well plates. After 24 hours, the cells were exposed to ionizing radiation (IR) at 4 Gy using an irradiator (MultiRad 160). Cells were harvested at different time after irradiation, washed twice with cold PBS, and fixed in 70% ethanol at –20°C for overnight. Fixed cells were washed with PBS and incubated with RNase A (100 µg/mL, 30 min, 37°C) and stained with propidium iodide (PI, 50 µg/mL) for 30 min at room temperature in the dark. The samples were analyzed by flow cytometry.

### 2.8. Isolation of subcellular fractions and chromatin-bound proteins

To isolate cytoplasmic, soluble nuclear, and chromatin-binding proteins, the cells were lysed in 500 µl ice-cold Buffer A [50 mM HEPES-KOH (pH 7.5), 140 mM NaCl, 1 mM EDTA (pH 8.0), 10% glycerol, 0.5% Nonidet P-40, 0.25% Triton X-100, 1 mM DTT, 1×protease inhibitor cocktail]. After centrifugation (700×g, 10 min), the supernatant was collected as the cytoplasm fraction and pellets were washed with Buffer A and resuspended in ice-cold Buffer B [10 mM Tris–HCl (pH 8.0), 200 mM NaCl, 1mM EDTA (pH 8.0), 0.5 mM EGTA (pH 8.0)], 1×protease inhibitor cocktail]. After extraction and centrifugation (20,000×g, 10 min), the soluble nuclear extract (NE) was collected. The pellet was washed with Buffer B and resuspended in Buffer C [500 mM Tris-HCl (pH 6.8), 500 mM NaCl, 1×protease inhibitor cocktail]. The suspension was sonicated for 2×10s and centrifuged (20,000×g, 10 min), and the supernatant was collected.

### 2.9. Immunofluorescence

Cells were seeded into 12-well plates with pre-placed coverslips at a density of 1×10^5^ cells/well and cultured overnight. The cells were pretreated with DMF for 3 h. Following HU treatments, the cells were fixed in ice-cold phosphate-buffered saline (PBS) with 0.1% Triton for 2 min, and then fixed in 4% paraformaldehyde. The cells were incubated with primary antibodies diluted in PBS (1:500) at 4°C overnight. After washing with PBS, Alexa-488-conjugated secondary antibodies (ThermoFisher Scientific) were applied for 2 h at room temperature. Nuclei were stained with DAPI (Sigma). Microscope images were captured using fluorescence microscopy (ZEISS Axio Observer).

### 2.10. HR-GFP reporter assay

U2OS cells containing a single copy of the HR repair reporter substrate DR-GFP were the gift from Dr. Jeremy Stark (City of Hope, Duarte, CA) [28]. U2OS cells were seeded in 12-well plates at a density of 2×10^5^ cells/well. After treatment with DMF or FH inhibitor for 3 h, the cells were infected with SceI adenovirus and replaced with fresh medium after 18h infection. The cells were collected after trypsin digestion, added with 200 µl 10% formaldehyde to 400 µl cell suspension, and kept in 4°C. Flow cytometry analysis was performed to quantify GFP positive cells.

### 2.11. Purification of recombinant proteins

To prepare purified wildtype and mutant RPA1 proteins, plasmids of Flag-RPA1 or Flag-RPA1 C481, C486 mutants were transiently transfected into HEK293T cells. After 48 h, the cell pellets were lysed with lysis buffer containing 50 mM Tris–HCl (pH 7.5), 150mM NaCl, 0.1%TritonX-100 and protease inhibitors. Lysates were sonicated and centrifuged at 14000g for 20min at 4°C. Supernatants were incubated with magnetic anti-flag beads overnight at 4°C with rotation. Beads were washed with lysis buffer for 5 times and eluted with 250 mg/ml Flag peptide (Sigma-Aldrich).

pET-28b-RPA1-WT were generated by insertion of the PCR fragment encoding RPA1 into the pET-28b vector. pET-28b-RPA1 plasmids were transformed into *E. coli* BL21 competent cells (New England Biolabs). 6xHis-tagged RPA1 proteins were expressed with 1 mM Isopropyl β-D-thiogalactopyranoside (IPTG) induction at 37°C for 4 hours and purified according to the protocol for purification of 6xHis-tag proteins. Briefly, cell pellets were lysed in buffer [50 mM Tris–HCl (pH 7.5), 500 mM NaCl, 5 mM imidazole, 1 mg/ml lysozyme, 1 mM PMSF, and protease inhibitor cocktails (Sigma)] on ice for 30 min, followed by sonication. The supernatant was collected by centrifugation at 11,000×*g* for 10 min at 4 °C.

### 2.12. Electrophoresis mobility shift assay (EMSA)

Purified RPA1 was incubated with sodium fumarate in 0.05M Tris-HCL (PH 7.5) buffer for 2h at 37°C. FAM-labeled single strand oligo (40nt) was heated at 95°C for 15 min and rapidly cooled down in ice-water. The indicated amount of RPA1 was incubated with 50nM oligo at room temperature for 30min in a reaction mixture consisting of 50 mM Tris (pH7.5), 5mM MgCl2, 50mM NaCl, 5mM EDTA and 0.1µg/µl BSA. Native PAGE gel (6%) was used to analyze the protein-DNA complexes with 100 volts for 45 min. The gels were imaged with a Typhoon FLA9500 scanner (GE Healthcare).

### 2.13. Statistical analysis

Statistical analyses were performed using GraphPad Prism 10.0. Data are expressed as mean ± SD. For comparison between two groups, an unpaired Student’s t-test was used for normally distributed data. Multiple group comparisons were conducted using one-way ANOVA followed by Tukey’s post hoc test. A *p*-value < 0.05 was considered statistically significant. *p* < 0.05 (*), *p* < 0.01 (**).

## 3. Results

### 3.1. Fumarate impairs CHK1, but not CHK2 checkpoint, activation in response to ionizing radiation (IR)

Cell survival relies on constant and strong genome surveillance systems that ensure the maintenance of genomic integrity and inheritance of genomic information. The ATM-CHK2 and ATR-CHK1 checkpoint signaling pathways are the main genome surveillance machinery in copying with endogenous and exogenous genotoxins [29]. To investigate whether oncometabolite fumarate affects DNA damage-induced activation of ATM-CHK2 and ATR-CHK1 checkpoint signaling, we treated both FH wild-type HK2 and 786-0 and FH mutant UOK262 and UOK268 cells with ionizing radiation (IR) and determined ATM-CHK2 and ATR-CHK1 checkpoint signaling activation by assessing the phosphorylation of CHK2 and CHK1, respectively. The results showed that phosphorylated CHK1and CHK2 were dramatically increased in response to IR in FH wild-type HK2 and 786-0 cells. CHK2 phosphorylation levels were also enhanced in FH mutant UOK262 and UOK268 cells. In sharp contrast, CHK1 activation was impaired in FH mutant cells, especially in UOK268 cells (Fig. 1A).

**Fig. 1.**
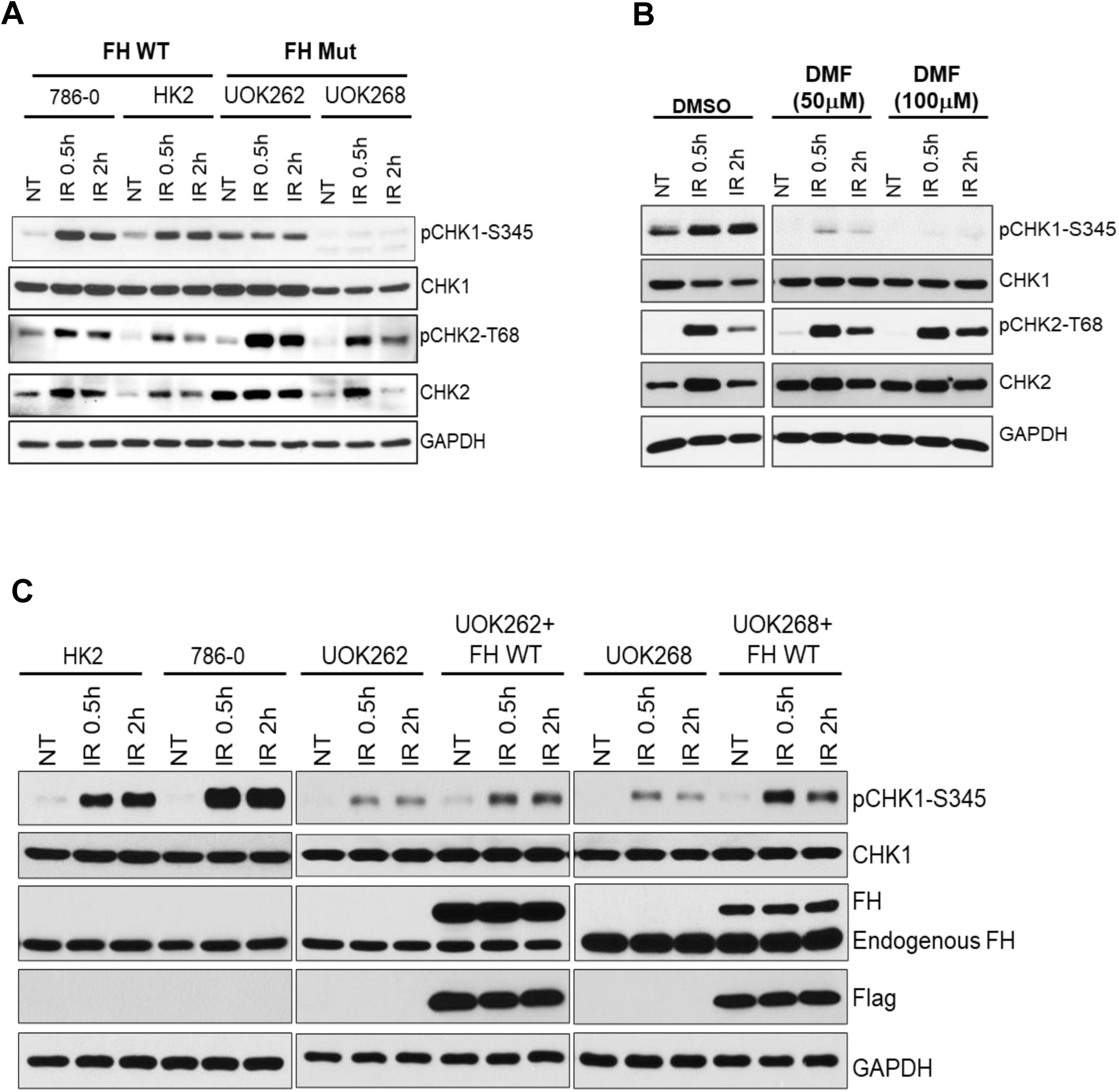
Fumarate accumulation impairs ATR-CHK1 but not ATM-CHK2 checkpoints activation in response to ionizing radiation (IR). A. FH wild-type and FH mutant cells were treated with IR. Cells were lysed for immunoblotting analyses. B. After treated with DMF (50µM or 100µM) for 3h, HK2 cells were exposed to IR. Cells were lysed for immunoblotting analyses. C. FH wild-type, FH mutant cells and FH mutant cells overexpression Flag-FH were treated with IR. Cell lysates were analyzed by immunoblotting.

To further confirm the results, we pretreated HK2 cells with dimethyl fumarate (DMF), which is a methyl ester of fumaric acid that is commonly used for mimicking biological function of fumarate. The activation of CHK1 was impaired obviously after DMF treatment in response to IR. Similar to FH mutant cells, DMF treatment had no effect on CHK2 phosphorylation (Fig. 1B). We next examined whether the CHK1 activation can be reversed when FH mutant UOK262 and UOK268 cells were re-introduced with wild type FH. The results showed the phosphorylation of CHK1 was dramatically increased in UOK262 and UOK268 cells with FH stable expression compared with the parental FH mutant cells (Fig. 1C). Collectively, these results showed that fumarate impaired CHK1 activation in response to IR.

### 3.2. Fumarate accumulation alleviates CHK1 activation in response to replication stress

Replication stress resulted from slowed DNA replication or stalled replication forks activate the ATR-CHK1 pathway to stabilize replication forks and arrest the cell cycle to allow for repair [30]. To further investigate whether fumarate affect CHK1 pathway in response to replication stress, we treated cells with hydroxyurea (HU). In HK2 and 786-0 cells, increased activation of CHK1 was observed but the activation is significantly attenuated in FH mutant cells. However, the attenuation is partially restored in the FH mutant cells overexpressing wild type FH (Fig. 2A). To further confirm the observation, both DMF and FH inhibitor were used. Consistently, CHK1 activation was dramatically decreased after DMF treatment in response to HU (Fig. 2B). FH inhibitor also reduced the activation of CHK1 by HU, especially in 48h treatment group (Fig. 2C). Taken together, these results demonstrated that the fumarate impaired CHK1 activation in response to replication stress as well.

**Fig. 2.**
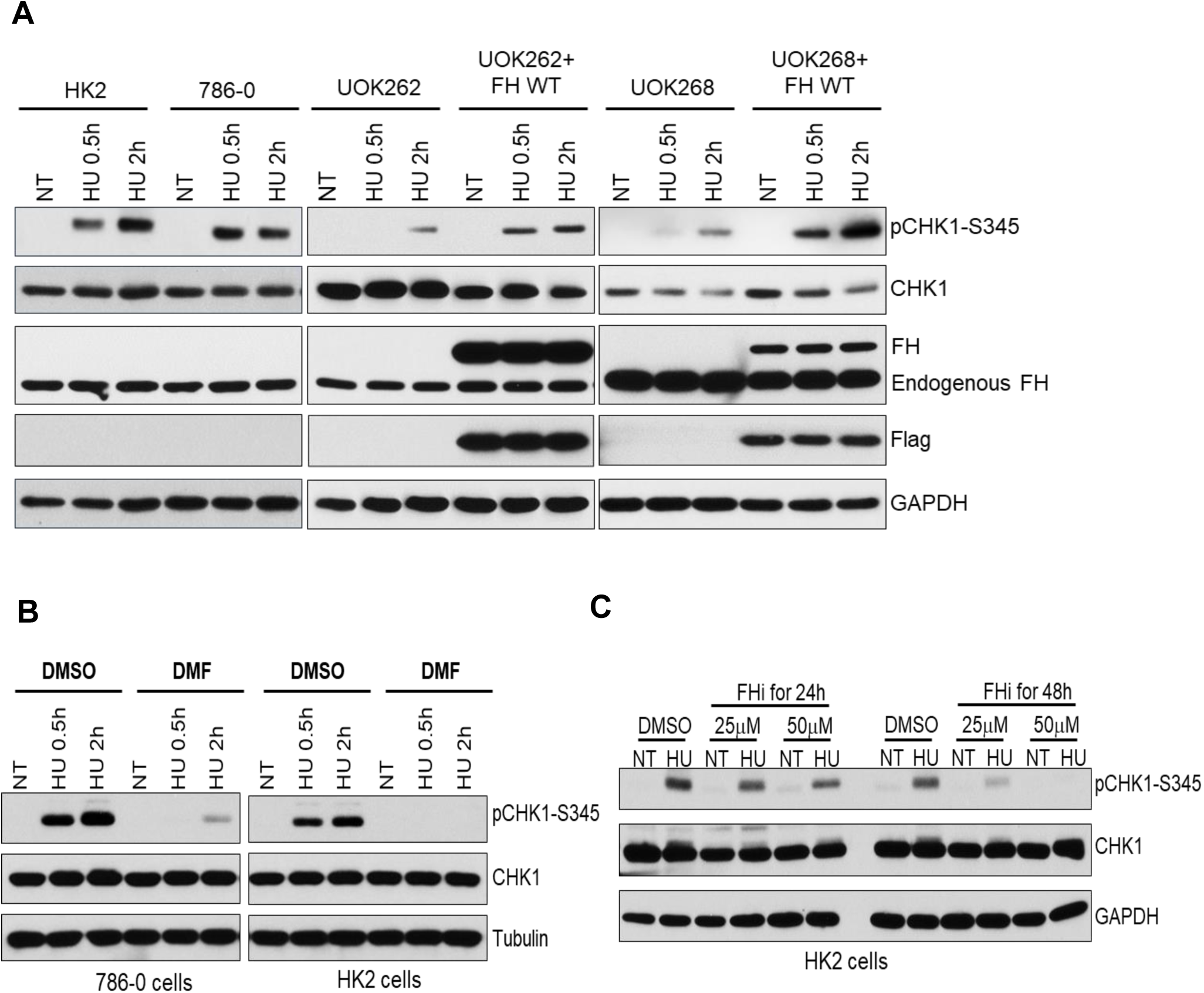
Fumarate accumulation impairs CHK1 activation in response to replication stress. A. FH wild-type, FH mutant cells and FH mutant cells overexpression Flag-FH were treated with HU. Cell lysates were analyzed by immunoblotting. B. After treated with DMF (100µM) for 3h, 786-0 and HK2 cells were exposed to IR. Cells were lysed for immunoblotting analyses. C. After incubated with FH inhibitor (25µM or 50µM) for 24h or 48h, HK2 cells were treated with HU. Cell lysates were analyzed by immunoblotting.

Activated CHK1 regulates the intra-S and G2/M checkpoints. To investigate the role of fumarate on cell cycle progression, cell cycle analysis was performed. FH normal, FH mutant, and FH mutant cells overexpressing wild type FH were treated with IR, respectively. In FH normal cells, there was a 2-fold increase in the percentage of cells in the G2/M phase after IR treatment (Fig. 3A and 3B). However, no increase in G2/M phase cell population was observed in mutant cells. The proportion of G2/M phase cells increased partially in FH mutant cells with ectopic wild type FH overexpression. These data indicated that FH mutation resulted in defects of G2/M phase arrest in response to IR which was accompanied by impairment of CHK1 activation.

**Fig. 3.**
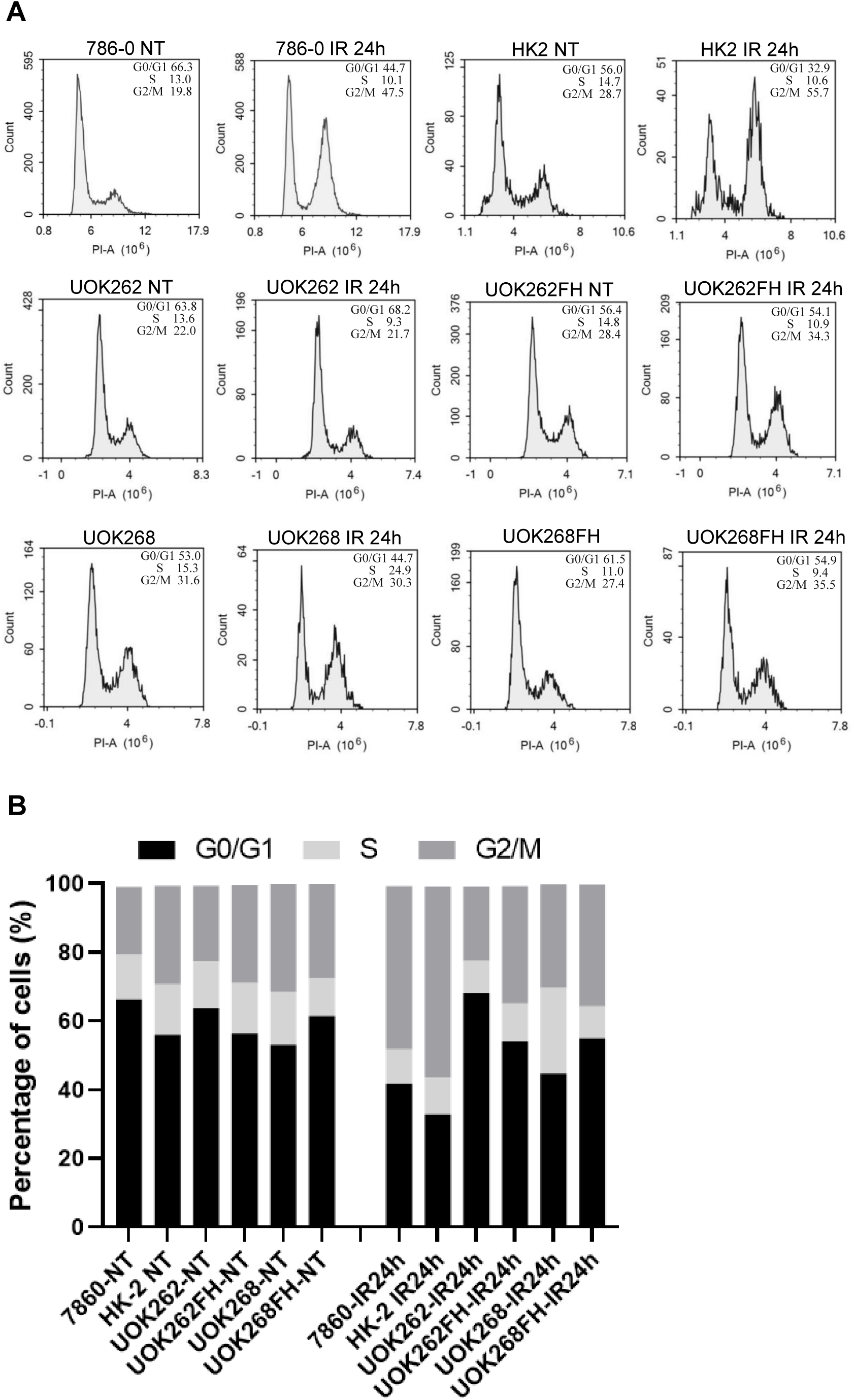
Fumarate hydrogenase (FH) mutation results in defects of G2/M phase arrest in response to IR. A. FH wild-type, FH mutant cells and FH mutant cells overexpression Flag-FH were treated with IR. Cell cycle analysis was performed using flow cytometry. The distributions of cells in different phases of the cell cycle are shown. B. Cell-cycle distribution is represented as a histogram.

### 3.3. Fumarate reduces homologous recombination (HR)

ATR-CHK1 activation enforces the S/G2 DNA damage checkpoint, delaying mitotic entry to allow time for DNA repair and maintenance of genomic integrity [31]. To examine the effect of fumarate accumulation on DNA damage repair, the DR-GFP reporter assay was performed to evaluate HR activity [28]. Consistent with the role for ATR-CHK1 in promoting HR, the efficiency of HR in the cells with DMF treatment was dramatically decreased in a doge dependent manner (Fig. 4A). In addition, similar results were observed in the cells treated with FH inhibitor (Fig. 4C). These results suggest that fumarate accumulation attenuates HR efficiency.

**Fig. 4.**
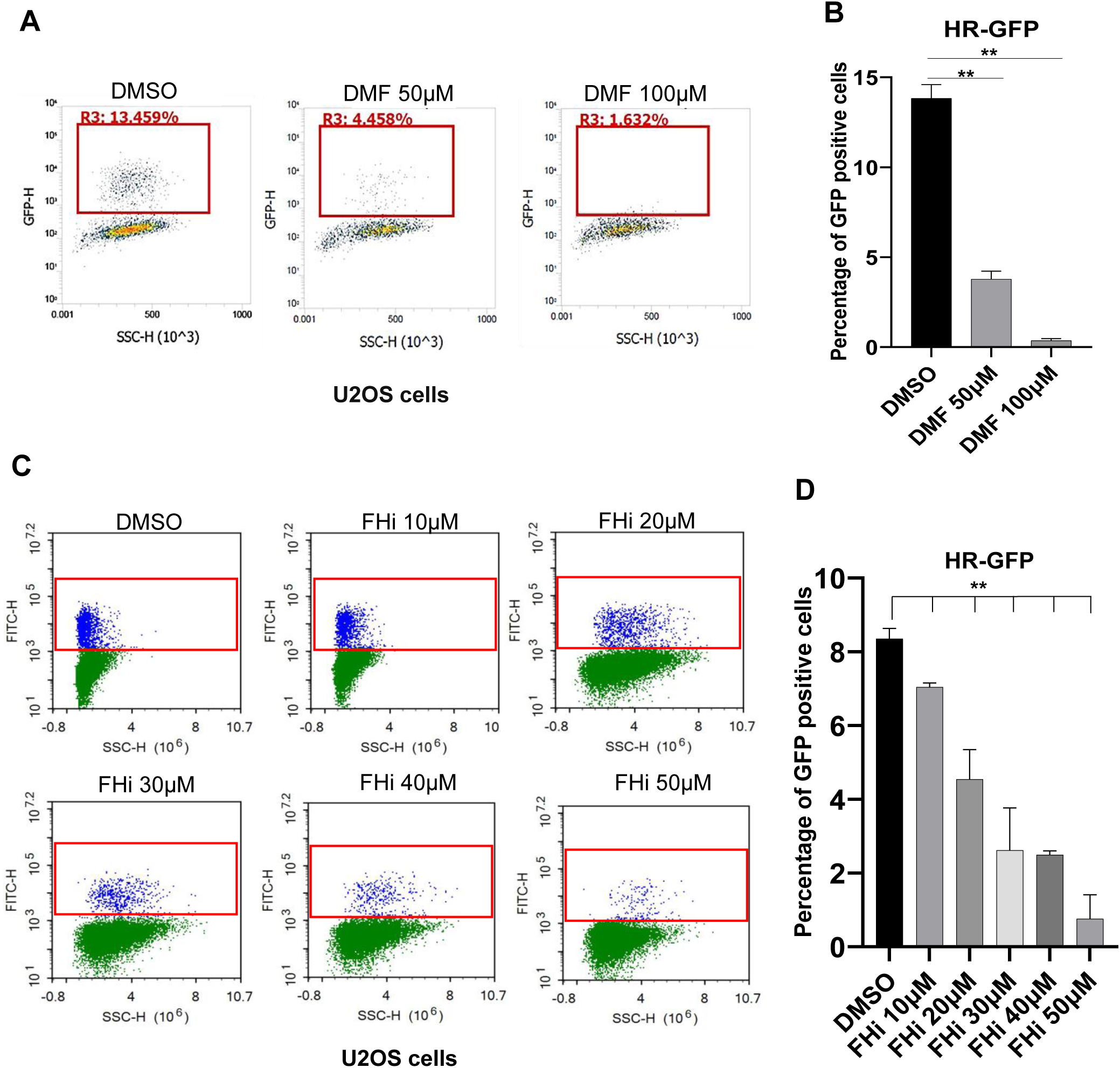
Dimethyl fumarate (DMF) and Fumarate hydrogenase inhibitor (FHi) reduce the cellular homologue directed recombination repair (HR) efficiency. A. UO2S cells were treated with DMF. Representative flow cytometry plots showing GFP-positive cells after I-SceI expression. Mean and standard deviation (SD) from three independent replicas is shown. B. UO2S cells were treated with FH inhibitor. Percent GFP positive cells were determined for HR efficiency. Data are presented as mean ± SD from three independent replicas.

### 3.4. Fumarate inhibits chromatin recruitment of ATR activation machinery in response to HU and IR

The ATR-CHK1 checkpoint is initiated by RPA-coated ssDNA, which recruits ATR via ATRIP, followed by further recruitment of TOPBP1 for efficient ATR-CHK1 axis activation [16]. To explore how fumarate affects ATR-CHK1 signaling pathway, chromatin-binding proteins were extracted to examine these signaling pathway proteins in HK2 cells treated with or without DMF. In the DMF treated cells, the chromatin-bound RPA1, ATRIP, ATR, and TopBP1 were markedly decreased compared with the untreated group in response to either HU or IR (Fig.5A). No significant change was observed in the cytoplasmic fractions (Fig.5B). Next, we examined RPA1 foci formation by immunofluorescence. The number of RPA1 foci was reduced in DMF-treated cells compared with the untreated group in response to HU (Fig. 5C and 5D). These data demonstrated that fumarate not only reduced the formation of RPA1 foci, but also impaired recruitment of checkpoint proteins to sites of DNA damages in response to DNA damages.

**Fig. 5.**
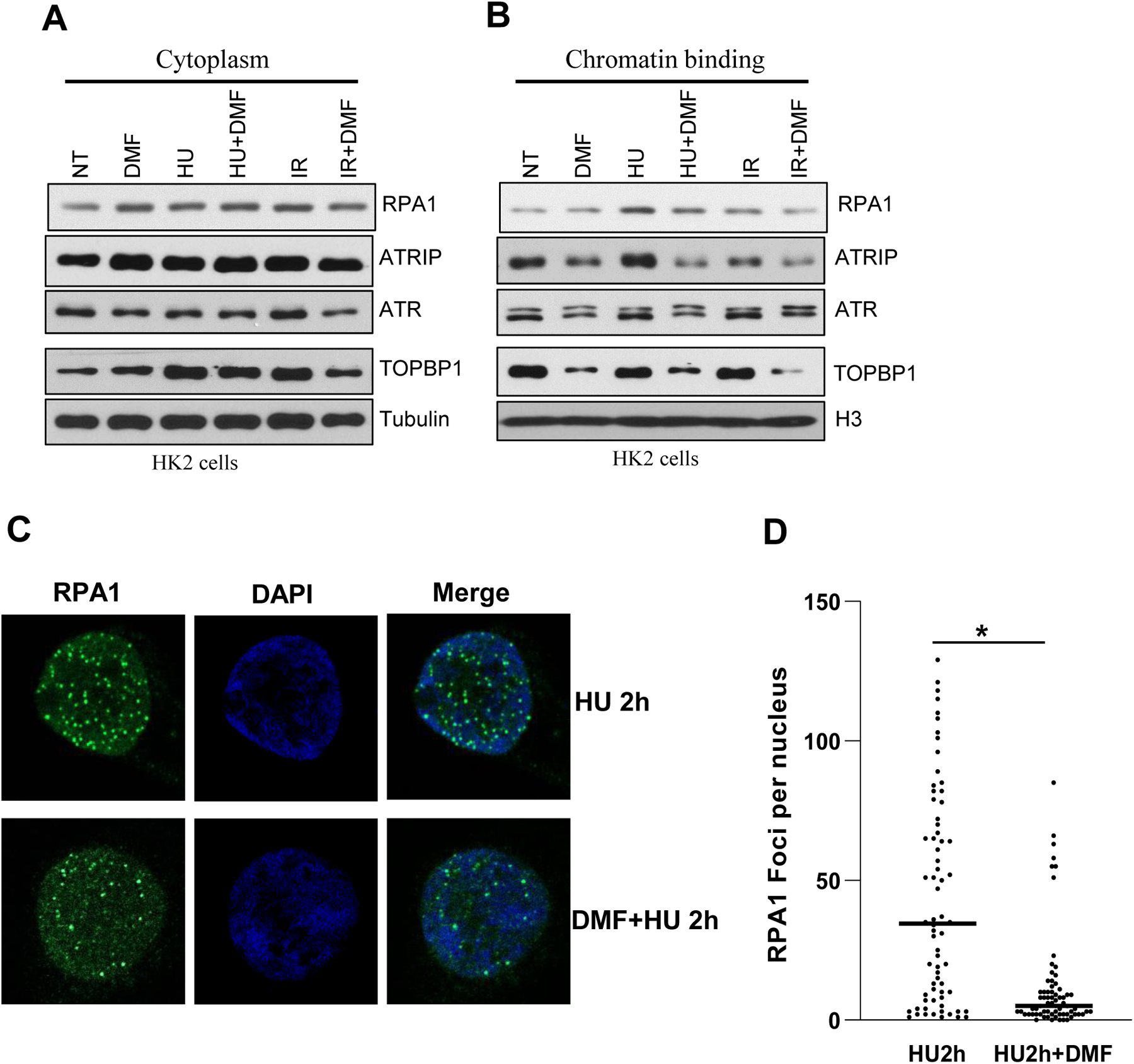
Dimethyl fumarate (DMF) inhibits chromatin recruitment of ATRIP-ATR in response to HU and IR treatments to cells. A,B. HK2 cells were pretreated with DMF (100µM) for 3h, followed by HU or IR treatment. Cells were fractionated into cytoplasmic and chromatin-bound fractions. Representative Western blots showing the levels of proteins in each fraction. C. HK2 cells were pretreated with DMF (100µM) for 3h, followed by HU treatment. Cells were stained with RPA1 antibody. D. Nuclear RPA1 foci were quantified using Image J.

### 3.5. RPA1 is modified by succination

Succination is a chemical reaction between fumarate and the thiol group of cysteine and form a stable thioether named 2-succino-cysteine (2SC) [32]. Protein succination can be detected by anti-2SC antibody. To investigate the mechanism by which fumarate impairs the chromatin recruitment of ATR-CHK activation machinery in response to HU and IR, protein co-immunoprecipitation and LC-MS/MS were performed to identify proteins that can be succianted by fumarate accumulation. Total cell lysates extracted from UOK268 cells were pulled down by anti-2SC antibody and subjected to LC-MS/MS analysis. A total of 930 proteins were identified (Table S1). Gene Ontology (GO) enrichment analysis based on MS data showed that most succinated proteins are involved in nucleic acids metabolism, including 25 ssDNA-binding proteins (Fig. 6A, Table S2). RPA1 was identified among these proteins. RPA1 is the essential protein of RPA complex that provides the platform to recruit ATR activation machinery for CHK1 activation under replication stress. Succination of RPA1 may be implicated in the impaired activation of CHK1. To test this, we examined whether RPA1 can be succinated *in vitro* by incubating purified RPA1 protein with sodium fumarate. The results showed that RPA1 could be succinated in a dose dependent manner (Fig. 6B). RPA consists of RPA1, RPA2, and RPA3. RPA1 and RPA2 play crucial roles in ssDNA binding and protein-protein interactions. We then examined whether RPA2 or RPA3 can also be succinated. RPA1, RPA2, and RPA3 were expressed and purified together as previously described [33]. Purified RPA complex was incubated with sodium fumarate, and protein succination was examined by anti-2SC antibody in immunoblotting. We found that only RPA1 was succinated, which was consistent with our Co-IP/LC-MS/MS data (Fig. 6C). To confirm that RPA1 is succinated inside cells, we overexpressed RPA1 in 293T cells treated with or without DMF, followed by immunoprecipitation by anti-Flag beads and immunoblotting. The results revealed that RPA1 was also succinated upon DMF stimulation in 293T cells (Fig. 6D). Collectively, these findings showed that RPA1 could be modified by succination, revealing an additional regulation of RPA1 at post-translational levels.

**Fig. 6.**
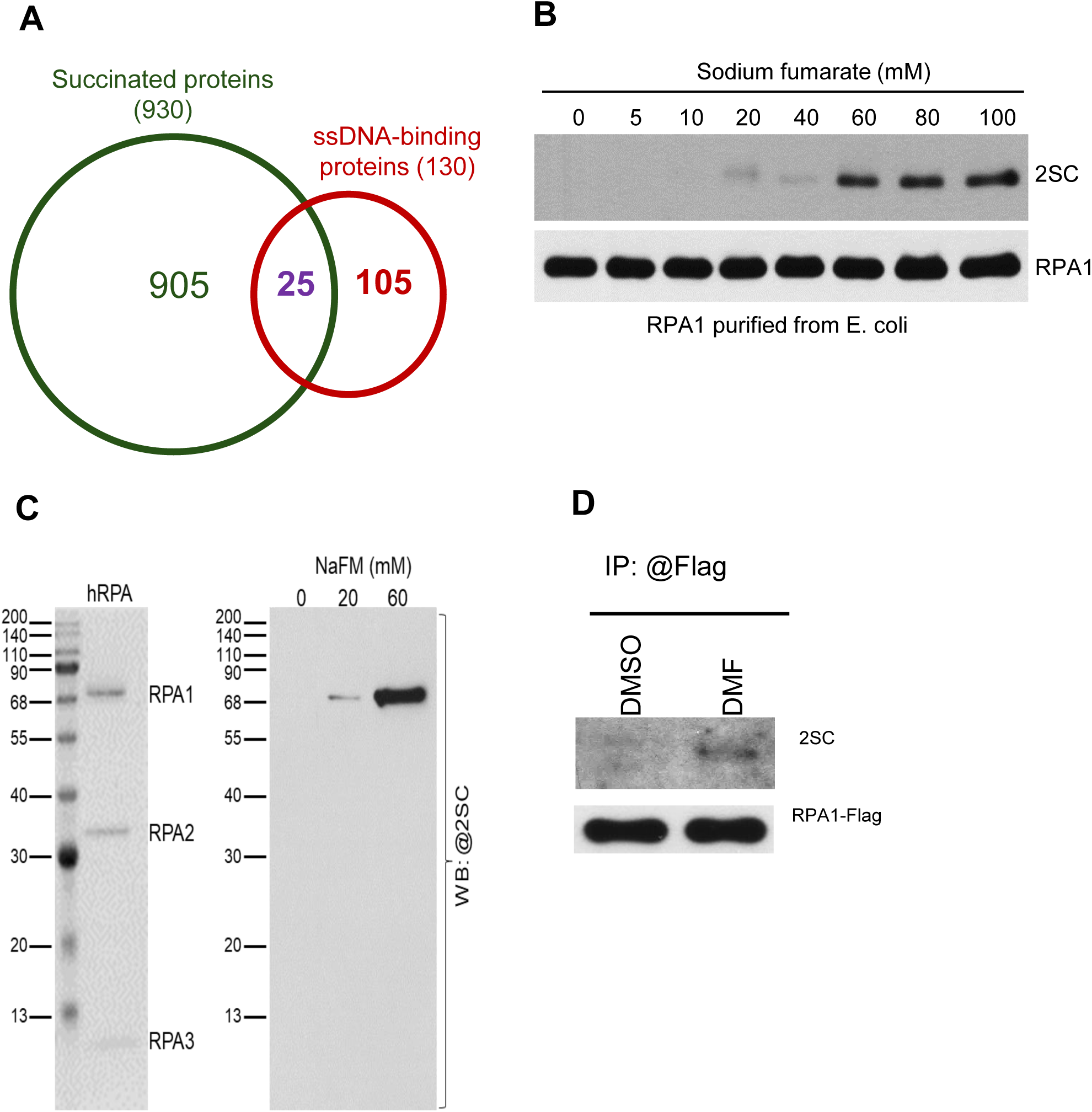
RPA1 is modified by succination. A. The proportion of ssDNA binding proteins among all succinated proteins based on LC-MS/MS analysis. B. Purified RPA1 was incubated with NaFM for 2h at 37°C, followed by immunoblotting analysis for the detection of 2-SC levels. C. Purified RPA complex, consisting of RPA1, RPA2, and RPA3, was incubated with NaFM for 2 h at 37 °C, followed by immunoblotting analysis to detect 2-SC levels. D. HEK293T cells were transfected with Flag-RPA1 and, 48 h later, treated with 100 µM DMF for 3 h. Cell lysates were immunoprecipitated with anti-Flag antibody, followed by immunoblotting analysis to detect 2-SC levels.

### 3.6. Fumarate promotes succination of RPA1 at zinc finger motif which affects its ssDNA-binding affinity

To identify the succinated cysteine residues in RPA1, purified protein was incubated with sodium fumarate, and then the succinated protein was analyzed by mass spectrometry (MS). MS results suggested that three cysteine residues of RPA1, cysteine 323 (C323), cysteine 481 (C481) and cysteine 486 (C486) were potentially succinated (Fig. 7A, B). To confirm the succinated sites, RPA1 wild-type and C481S, C486S double mutant proteins were purified from 293T cells (Fig. 7C). *In vitro* succination assay showed the succination of RPA1 obviously decreased with C481S, 486S mutants (Fig. 7D). Cysteine residues C481 and C486 in RPA1 are located within a zinc finger-like motif which is critical for stabilizing the structure of the DNA-binding domains of RPA1. Alteration of structure of RPA1 reduces its ssDNA-binding activity [34]. To confirm whether RPA1 binding was influenced by succination, we examined the ssDNA-binding capacity of modified RPA1 by electrophoretic mobility shift assay (EMSA). RPA1 protein bound ssDNA-binding activity with an increasing amount of RPA1. By contrast, ssDNA-binding activity of succinated RPA1 was significantly reduced (Fig. 7E). These data collectively suggest that succination of RPA1 in sites of C481 and C486 destroys the zinc finger-like motif and reduces its ssDNA-binding activity.

**Fig. 7.**
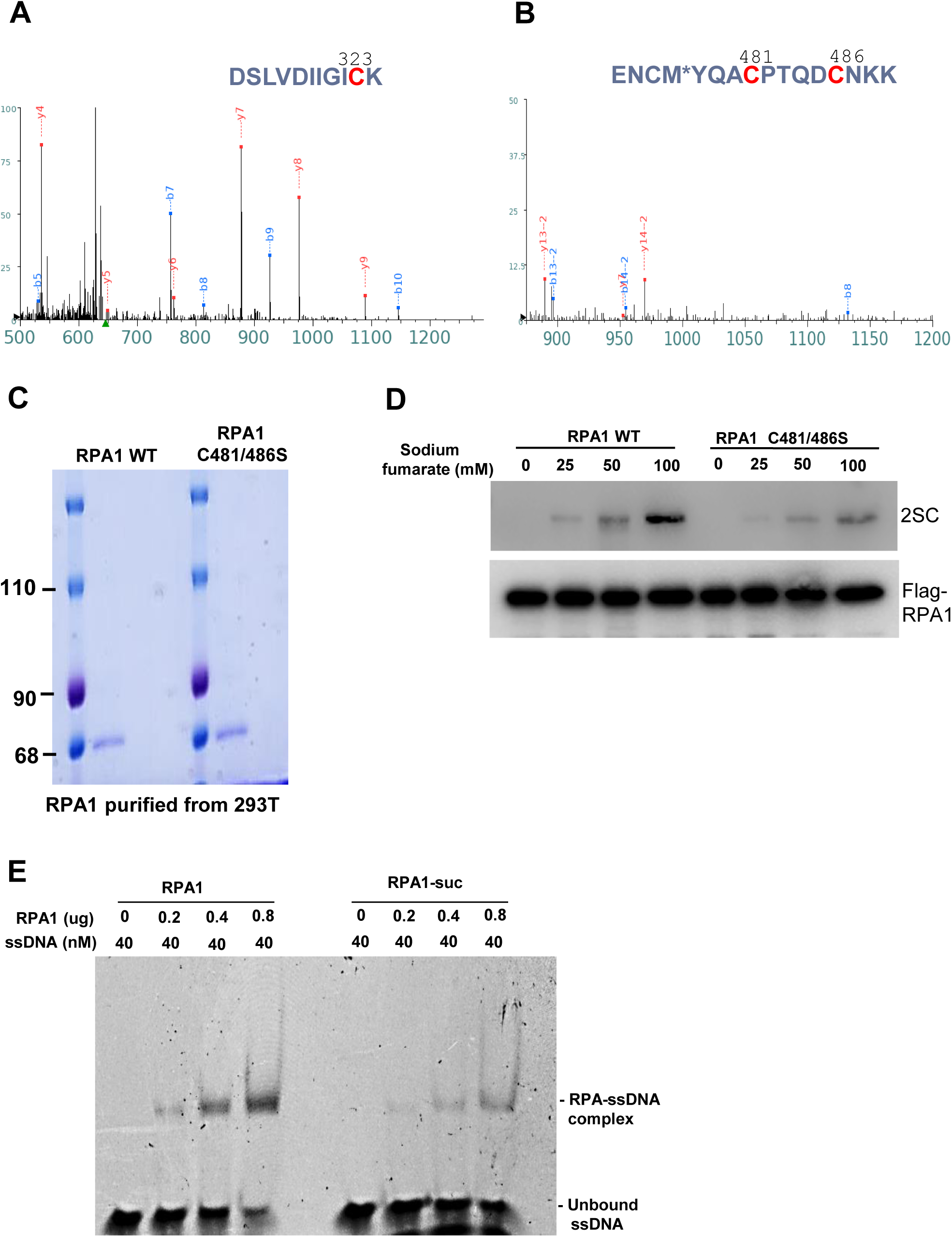
Fumarate promotes succination of RPA1 in Zinc finger motif, which affects its ssDNA-binding activity. A, B. Purified recombinant RPA1 protein was incubated with NaFM and subjected to SDS-PAGE. Proteins were visualized by Coomassie staining, and the gel bands corresponding to RPA1 were excised and analyzed by LC-MS/MS. Mass spectrometry identified 2-SC modifications at cysteine residues of C323, C481 and C486. C. Purified recombinant RPA1 proteins, including wild-type and C481/486 mutant, were analyzed by SDS-PAGE and visualized by Coomassie staining. Both proteins showed predominant bands at the expected molecular weight (70kDa). D. Purified recombinant RPA1 proteins, including wild-type (WT) and C481/486 mutant were was incubated with NaFM for 2 h at 37 °C, followed by immunoblotting analysis to detect 2-SC levels. E. EMSA was performed to examine the binding of RPA1 to ssDNA. Purified recombinant proteins were incubated with a FAM-labeled ssDNA, and the complexes were subjected to a native polyacrylamide gel. RPA1-ssDNA complexes appeared as the top band on the gel. The lower bands represent the free ssDNA probe.

## 4. Discussion

In this study, we show that fumarate accumulation leads to succination of RPA1 at zinc finger motif cysteine residues 481 and 486, which reduces its ssDNA-binding affinity. Consistently, fumarate inhibits chromatin recruitment of ATR activation machinery and hence abolishes CHK1 activation in response to DNA damage and replication stress. Consequently, fumarate reduces homologous recombination and G2/M phase arrest in response to DNA damage. Our results reveal a novel mechanism by which metabolic reprogramming contributes to genomic instability in cancer cells. Our findings may provide alternative therapy strategies for cancers with mutations of the key enzymes in TCA cycle, especially FH-dRCC.

FH is a nuclear-encoded metabolic enzyme and a key component of the TCA cycle that catalyzes the reversible conversion of fumarate to malate in mitochondria. In the cytosol, FH metabolizes fumarate, which is a byproduct of the urea cycle and amino acid catabolism [35]. FH translocates from the cytosol into the nucleus upon DNA damage and participates in the DNA damage response (DDR). For example, yeast cells lacking cytosolic FH exhibit increased sensitivity to DSB inducing agents. It has been reported that FH facilitates efficient HR [36]. It was reported that FH loss and fumarate accumulation lead to a weakened G2 checkpoint, predisposing cells to endogenous DNA damage while simultaneously conferring resistance to IR [37]. These facts suggest that FH is not only a metabolic enzyme but also a guardian of genome stability, linking metabolism to DNA repair [38]. In this study, we demonstrate that loss of FH not only impairs the CHK1 checkpoint but also disrupts HR. Our results have identified a novel mechanism by which fumarate accumulation in FH-deficient cells contributes to genomic instability and hence tumorigenesis.

RPA1 plays a central role in DNA replication, repair, and the activation of the ATR-CHK1 checkpoint pathway in response to replication stress or DNA damage [39, 40]. It is a critical ssDNA-binding protein required for ATR-CHK1 pathway activation. RPA1 contains a zinc finger motif, which is essential for its high-affinity binding to ssDNA. We demonstrate that RPA1 undergoes succination at cysteine residues 323, 481, and 486, with the latter two sites located within the zinc finger motif of RPA1. Succination at these critical cysteines may disrupt the structural integrity of the zinc finger, reducing the binding affinity of RPA1 to ssDNA, thereby impairing recruitment of TOPBP1 and subsequent CHK1 activation. Consistently, we find that FH wild type cells treated with DMF and FH mutant cells exhibited reduced ATR-CHK1 but not ATM-CHK2 checkpoint activation in response to either IR or HU, which was accompanied by defective G2/M arrest. Consistent with the important role for ATR-CHK1 in enhancing HR, we found FH inhibitor and DMF compromised HR repair capability. These findings suggest that impairment of ATR-CHK1 signaling may be the main mechanism of genome instability and hence aggressive tumor growth of FH-dRCC. Nevertheless, it will be important to determine how succination of cysteine residues 481 and 486 alter RPA1 structure itself and the RPA complex.

While fumarate-induced succination has been described in metabolic enzymes and transcription factors, our study is the first to demonstrate succination of RPA1 as a mechanism for defective checkpoint signaling. By connecting metabolic dysregulation to the ATR-CHK1 DNA damage response, our findings highlight how FH loss promotes genome instability not only through epigenetic alterations but also via direct modification of DNA repair machinery. This mechanistic insight provides a new perspective on the tumorigenic potential of FHdRCC cells. However, it will be intriguing to assess whether RPA1 succination precludes other RPA1 modifications which are essential for RPA to recruit ATRIP, TOPBP1 and others for effective ATR-CHK1 checkpoint activation in response to replication stress and DNA damages.

## 5. Conclusion

In summary, we discovered that abnormal oncometabolite fumarate accumulation or FH mutation leads to succination of many proteins important in nucleic acid metabolism. Particularly, we identified and validated that RPA1 is modified by succination, a post-translational protein modification resulting from dysregulation of the TCA cycle. Importantly, we found that RPA1 succination reduces its ssDNA binding affinity, which is accompanied by impaired ATR-CHK1 activation, defective G2/M arrest, and reduced HR in response to DNA damages. These findings have established a direct link between the oncometabolite fumarate and impaired DNA damage response in FH-deficient renal cell carcinoma (FH-dRCC).

Supplementary Materials can be found on the *Cancer Letter* website.

## Acknowledgements

The authors wish to thank Dr. Feiyan Pan and Zhigang Guo in Nanjing Normal University for their support in the initial stage of the project. We thank W. M. Linehan at NCI Urologic Oncology Branch for his unique cancer cell lines, UOK262 and UOK268 and L. Brown and B. Armstrong at the COH Flow Cytometry and Light Microscopy core facilities for their technical support. This work was supported by NIH grants R01 CA085344 to B.H.S and R50 CA211397 to L.Z. Research reported in this publication includes work performed using City of Hope shared resources supported by the National Cancer Institute of the National Institutes of Health under award number P30 CA033572.

## Authors’ Contributions

**N. Liu:** Conceptualization, Data curation, Formal analysis, Investigation, Methodology, Writing – original Draft. **C. Shen:** Conceptualization, Data curation, Formal analysis, Investigation, Methodology, Validation, Writing – review & editing. **T. Zhou:** Conceptualization, Data curation, Investigation, Methodology. **Y. Zhou:** Data curation, Investigation, Methodology. **H. Yuan:** Data curation, Investigation. **H. Bi:** Investigation, Resources. **Y. Wang:** Data curation. **X. Pei:** Investigation. **R. Su**: Data curation. **L. Zheng:** Funding acquisition, Writing – review & editing. **Y. Shi:** Project administration, Resources, Writing – review & editing. **B. Shen:** Conceptualization, Data curation, Project administration, Supervision, Funding acquisition, Writing – review & editing.

## Supplementary Information

**Supplementary Table-S1**: Top 30 PSMs among 930 succinated proteins identified by MS in UOK268 cells

**Supplementary Table-S2:** Succinated 25 ssDNA binding proteins identified by MS in UOK268 cells

