## Supplemental Table1 and Table 2 for "Oncometabolite fumarate impairs ATR-CHK1 signaling by succinating RPA1 in Fumarate Hydratase-deficient renal cell carcinoma cells"

**Supplementary Table S1: Top 30 PSMs among 930 succinated proteins identified by MS in UOK268 cells**

| Accession | GenSymbol | Protein names | AAs* | Summed PSM** |
|---|---|---|---|---|
| Q09666 | AHNK | Neuroblast differentiation-associated protein AHNAK | 5890 | 400 |
| P35580 | MYH10 | Myosin-10 (Cellular myosin heavy chain, type B) | 1976 | 198 |
| P07437 | TBB5 | Tubulin beta chain (Tubulin beta-5 chain) | 444 | 170 |
| P68371 | TBB4B | Tubulin beta-4B chain (Tubulin beta-2 chain) | 445 | 154 |
| P35579 | MYH9 | Myosin-9 (Cellular myosin heavy chain, type A) | 1960 | 147 |
| Q13885 | TBB2A | Tubulin beta-2A chain (Tubulin beta class IIa) | 445 | 138 |
| Q9BVA1 | TBB2B | Tubulin beta-2B chain | 445 | 136 |
| Q9BQE3 | TBA1C | Tubulin alpha-1C chain (Alpha-tubulin 6) | 449 | 128 |
| Q71U36 | TBA1A | Tubulin alpha-1A chain (Alpha-tubulin 3) | 451 | 135 |
| P04350 | TBB4A | Tubulin beta-4A chain (Tubulin 5 beta) (Tubulin beta-4 chain) | 444 | 126 |
| Q02413 | DSG1 | Desmoglein-1 (Cadherin family member 4) | 1049 | 121 |
| Q07954 | LRP1 | Prolow-density lipoprotein receptor-related protein 1 (LRP-1) | 4544 | 112 |
| P52272 | HNRPM | Heterogeneous nuclear ribonucleoprotein M (hnRNP M) | 730 | 106 |
| P0DOX5 | IGG1 | Immunoglobulin gamma-1 heavy chain | 449 | 107 |
| Q9UQE7 | SMC3 | Structural maintenance of chromosomes protein | 1217 | 91 |
| P19338 | NUCL | Nucleolin (Protein C23) | 710 | 90 |
| P60709 | ACTB | Actin, cytoplasmic 1 (Beta-actin) | 375 | 98 |
| P07355 | ANXA2 | Annexin A2 (Annexin II) (Annexin-2) (Calpactin I heavy chain) | 339 | 80 |
| P0DMV9 | HS71B | Heat shock 70 kDa protein 1B (Heat shock 70 kDa protein 2) | 641 | 76 |
| Q08211 | DHX9 | ATP-dependent RNA helicase A (EC 3.6.4.13 | 1270 | 75 |
| P06748 | NPM | Nucleophosmin (NPM) (Nucleolar phosphoprotein B23) | 294 | 74 |
| Q14683 | SMC1A | Structural maintenance of chromosomes protein 1A | 1233 | 74 |
| Q9BUF5 | TBB6 | Tubulin beta-6 chain (Tubulin beta class V) | 446 | 73 |
| Q13835 | PKP1 | Plakophilin-1 (Band 6 protein) (B6P) | 747 | 66 |
| Q14562 | DHX8 | ATP-dependent RNA helicase DHX8 (EC 3.6.4.13) | 1220 | 64 |
| P09874 | PARP1 | Poly [ADP-ribose] polymerase 1 (PARP-1) (EC 2.4.2.30) | 1014 | 64 |
| P23246 | SFPQ | Splicing factor, proline- and glutamine-rich | 707 | 64 |
| P01859 | IGHG2 | Immunoglobulin heavy constant gamma 2 | 326 | 110 |
| Q15233 | NONO | Non-POU domain-containing octamer-binding protein | 471 | 59 |
| P05388 | RLA0 | 60S acidic ribosomal protein P0 (60S ribosomal protein L10E) | 317 | 59 |

***AAs : amino acid numbers;  **PSM : peptide-spectrum matches**

**Supplementary Table -S2 :Succinated 25 ssDNA binding proteins identified by MS in UOK268 cells**

| Gene name | Protein name | PSMs |
|---|---|---|
| DHX9 | DEAH box protein 9 | 75 |
| HNRNPA2B1 | Heterogeneous nuclear ribonucleoproteins A2/B1 | 43 |
| HNRNPA1 | Heterogeneous nuclear ribonucleoprotein A1 | 42 |
| HNRNPU | Heterogeneous nuclear ribonucleoprotein U | 35 |
| SMC2 | Structural maintenance of chromosomes protein 2 | 14 |
| SMC4 | Structural maintenance of chromosomes protein 4 | 10 |
| POLR2H | DNA-directed RNA polymerases I, II, and III subunit RPABC3 | 7 |
| MCM7 | DNA replication licensing factor MCM7 | 7 |
| YBX1 | Y-box-binding protein 1 | 6 |
| HNRNPDL | Heterogeneous nuclear ribonucleoprotein D-like | 6 |
| SAMHD1 | Deoxynucleoside triphosphate triphosphohydrolase | 5 |
| SUB1 | Activated RNA polymerase II transcriptional coactivator p15 | 4 |
| RPA1 | Replication protein A 70 kDa DNA-binding subunit | 4 |
| HSPD1 | 60 kDa heat shock protein, mitochondrial | 4 |
| PCBP1 | Poly(rC)-binding protein 1 | 4 |
| PCBP2 | Poly(rC)-binding protein | 4 |
| NME1 | Nucleoside diphosphate kinase A | 4 |
| SSBP1 | Single-stranded DNA-binding protein, mitochondrial | 4 |
| POLR2G | DNA-directed RNA polymerase II subunit RPB7 | 3 |
| PURA | Purine-rich single-stranded DNA-binding protein alpha | 3 |
| TOP1 | DNA topoisomerase 1 | 3 |
| PURB | Purine-rich element-binding protein B | 2 |
| MCM2 | DNA replication licensing factor MCM2 | 1 |
| TDP2 | Tyrosyl-DNA phosphodiesterase 2 | 1 |
| REXO4 | RNA exonuclease 4 | 1 |

**Proteins marked in red are DNA binding only**
